# Armadillo repeat-containing protein 1 is a dual localization protein associated with mitochondrial intermembrane space bridging complex

**DOI:** 10.1101/656934

**Authors:** Fabienne Wagner, Tobias C. Kunz, Suvagata R. Chowdhury, Bernd Thiede, Martin Fraunholz, Debora Eger, Vera Kozjak-Pavlovic

## Abstract

Cristae architecture is important for the function of mitochondria, the organelles that play the central role in many cellular processes. The mitochondrial contact site and cristae organizing system (MICOS) together with the sorting and assembly machinery (SAM) forms the mitochondrial intermembrane space bridging complex (MIB), a large protein complex present in mammalian mitochondria that partakes in the formation and maintenance of cristae. We report here a new subunit of the mammalian MICOS/MIB complex, an armadillo repeat-containing protein 1 (ArmC1). ArmC1 localizes both to cytosol and mitochondria, where it associates with the outer mitochondrial membrane through its carboxy-terminus. ArmC1 interacts with other constituents of the MICOS/MIB complex and its amounts are reduced upon MICOS/MIB complex depletion. Mitochondria lacking ArmC1 do not show defects in cristae structure, respiration or protein content, but appear fragmented and with reduced motility. ArmC1 represents therefore a peripheral MICOS/MIB component that appears to play a role in mitochondrial distribution in the cell.

## Introduction

Mitochondria are dynamic organelles performing various important cellular functions. ATP production, β-oxidation of fatty acids and synthesis of iron-sulfur clusters all take place within the mitochondria. Furthermore, mitochondria have a central role in calcium homeostasis and programmed cell death [1].

Mitochondria are surrounded by two membranes, the outer (OMM) and the inner mitochondrial membrane (IMM). The OMM separates mitochondria from the cytosol and harbors porins that enable solute and small molecules exchange, as well as the translocase of the outer membrane (TOM), which serves as an entry point for all proteins transported into mitochondria. In addition, sorting and assembly machinery (SAM) in the OMM cooperates with the TOM complex to mediate the outer membrane integration of mitochondrial β-barrel proteins, which include previously mentioned porins [2]. The most abundant proteins of the IMM are those involved in the ATP synthesis by oxidative phosphorylation (OXPHOS). The IMM also contains translocase complexes, involved in protein import and sorting, as well as carrier proteins, necessary for metabolite exchange [3]. The IMM can be divided into the inner boundary membrane (IBM) and cristae region, which differ in protein composition. Cristae represent folds of the IMM that enlarge its surface and are connected to the IBM by tubular cristae junctions (CJ) [4]. The two membranes enclose two additional compartments of the mitochondria, the matrix, surrounded by the IMM, and the intermembrane space (IMS) between the OMM and the IMM.

Formation and maintenance of cristae and CJs has in the recent years been linked to the existence of the mitochondrial contact site and cristae organizing system (MICOS) [5–8]. In mammalian mitochondria, MICOS closely interacts with the SAM complex and forms the 2.2 – 2.8 MDa large mitochondrial intermembrane space bridging complex (MIB) [9–11]. The central component of the MICOS and MIB complex, Mic60/Mitofilin, is a protein anchored with an amino (N)-terminal anchor in the IMM [12, 13]. Depletion of Mic60/Mitofilin leads to a reduction in the amounts of other MICOS and MIB components [10], which can be used to identify proteins associated with the MIB complex [14]. Other IMM-localized MICOS components include several coiled-coil helix-coiled-coil helix domain (CHCHD)-containing proteins, as well as apolipoproteins and scaffolding proteins, of which Mic19/CHCHD3 and Mic10/MINOS1 appear to play the most important role for the cristae formation [11]. The OMM constituents of the MIB complex include the members of the SAM complex, Sam50 and Metaxins, as well as a member of the J protein family, DnaJC11 [15, 16].

In 2015, an in depth proteomic immunoprecipitation-mass spectrometry analysis of the MICOS complex was performed by Guarani and colleagues. As a result, several novel proteins associated with the MICOS/MIB complex were identified. Among these putative MICOS/MIB complex interactors was an armadillo repeat-containing protein 1 (ArmC1), which was co-precipitated with Mic13/Qil1, Mic27/ApoOL, Mic19/CHCHD3, Metaxin 2 and DnaJC11[17]. In this study, we have further characterized ArmC1 protein. We show that ArmC1 localizes to the cytosol, as well as to mitochondria. ArmC1 associates with, but is not integrated into the OMM. We confirm the data of Guarani and colleagues, showing that ArmC1 is indeed a component of the MICOS/MIB complex. Knockout of ArmC1, however, has no greater effect on mitochondrial respiration or the stability of the MICOS/MIB complex and cristae formation, even though mitochondria in cells lacking ArmC1 are slightly fragmented and with impaired motility. ArmC1, therefore, represents a novel peripheral component of the MICOS/MIB complex with an unusual localization to both cytosol and mitochondria, which might play a role in cellular distribution of mitochondria.

## Results

### ArmC1 is a protein with dual localization associated with mitochondrial outer membrane

As a part of the previous study, we have performed stable isotope labeling with amino acids in cell culture (SILAC) in combination with quantitative mass spectrometry to analyze the levels of mitochondrial proteins after depletion of Mic60/Mitofilin [14]. One of the significantly reduced proteins was ArmC1, a protein containing an armadillo repeat, which is a ~42 amino acid (aa) motif of three α-helices first characterized in the *Drosophila* segment polarity protein Armadillo [18]. In the meantime, ArmC1 was reported as a putative interactor of the MICOS/MIB complex [17]. Therefore, we endeavored to further characterize this protein.

We have cloned the gene for the longer isoform 1 of ArmC1 into a mammalian expression vector pCDNA3, introducing the FLAG-tag at either the N-terminus or the carboxy (C)-terminus of the protein (FLAG-ArmC1 and ArmC1-FLAG, respectively). Upon expression of these constructs in HeLa cells, we observed that whereas ArmC1-FLAG did not appear to specifically co-localize with mitochondria, but was evenly distributed in the cytosol, FLAG-ArmC1 was found largely associated with MitoTracker-stained mitochondria, although a smaller amount of protein was detected in the cytosol, as well. We repeated the experiments, this time introducing the green fluorescent protein (GFP) to either N- or C-terminus of ArmC1. The results were comparable, with ArmC1-GFP fluorescence distributed in the cytosol and GFP-ArmC1 signal mostly co-localizing with mitochondrial Tom20 signal, in addition to the weak cytosolic presence (Fig 1A). We analyzed GFP-ArmC1 mitochondrial localization in more detail using structured illumination microscopy (SIM). The GFP-ArmC1 signal in mitochondria co-localized with the OMM staining with Tom20, indicating surface localization of the protein. In addition, we noticed as before cytosolic presence of the GFP-fusion protein (Fig 1B). The expression of GFP-ArmC1 in cells led to fragmentation, aggregation and enlargement of mitochondria, but we did not observe this effect when we expressed FLAG-ArmC1 (Fig 1A and B).

**Fig 1.**
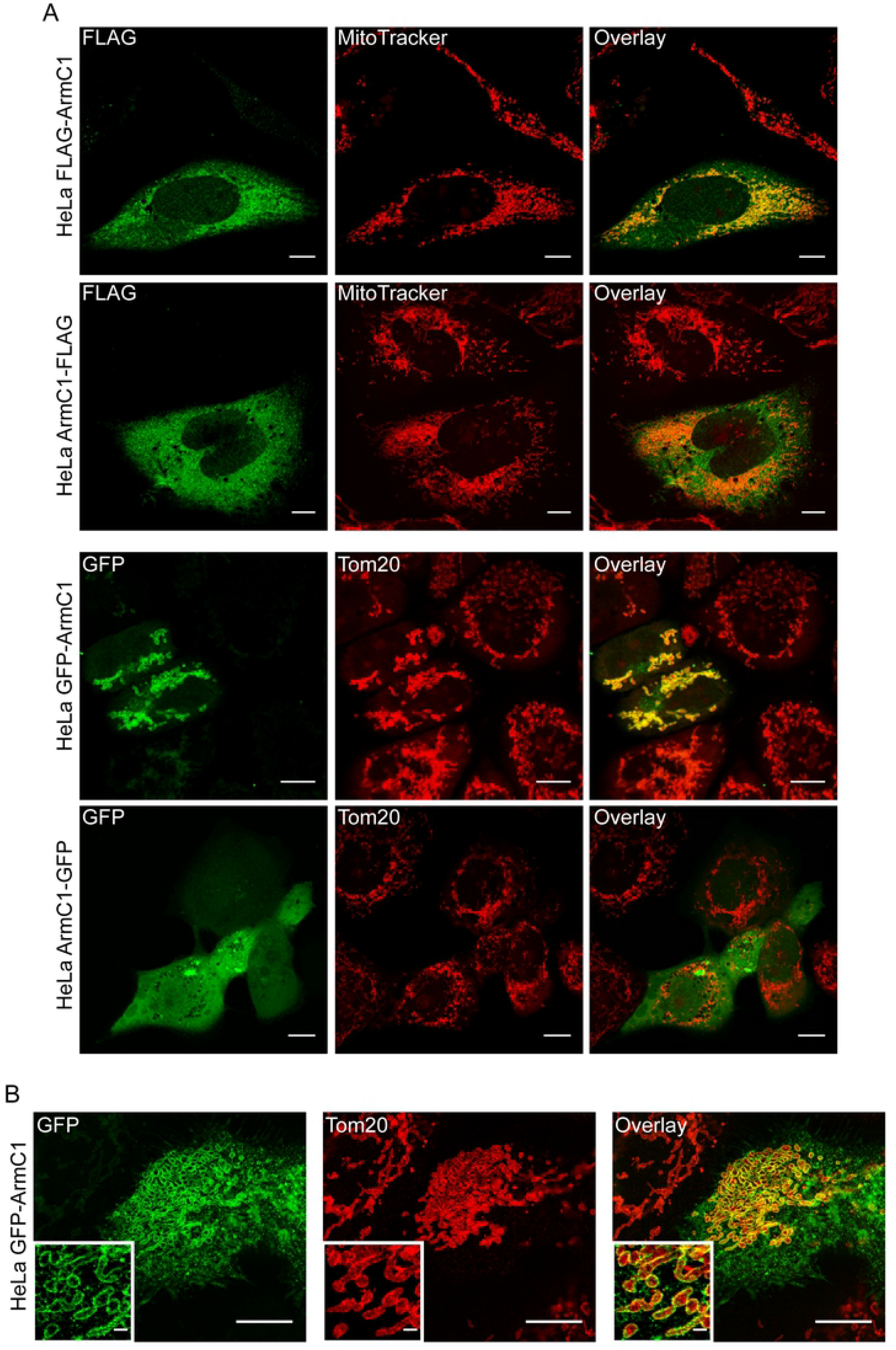
Localization of expressed ArmC1 in HeLa cells. (A) HeLa cells were seeded on coverslips and transfected with pCDNA3 plasmid carrying the information for the ArmC1 fused to the FLAG-tag or green fluorescent protein (GFP) either at the N-terminus (FLAG-ArmC1 and GFP-ArmC1) or at the C-terminus (ArmC1-FLAG and ArmC1-GFP). 24 h after transfection, mitochondria were stained with MitoTracker, followed by fixation and staining with anti-FLAG and secondary Cy2-coupled antibody for the FLAG-tagged constructs. In the case of the GFP-fused constructs, cells were fixed and stained with anti-Tom20 antibody, followed by Cy3-coupled secondary antibody. Samples were analyzed by confocal microscopy. Scale bar is 10 µm. (B) Cells in which GFP-ArmC1 construct has been expressed were prepared as in A and analyzed by structured illumination microscopy (SIM). Insets show magnified areas of the pictures. Scale bar is 10 µm, inset scale bar is 1 µm. Tom20 – translocase of the outer membrane 20.

To determine the cellular and sub-mitochondrial localization of ArmC1 we generated an antibody against ArmC1 and performed fractionation experiments using non-transfected HEK293T cells, as well as HEK293T cells that were transfected with pCDNA3 FLAG-ArmC1 construct. We separated the cytosolic from the heavy membrane fraction, which contained mitochondria. Western blot analysis showed that FLAG-ArmC1 was present in both mitochondrial and cytosolic fraction. The same was the case with the endogenous ArmC1 (Fig 2A). Protease treatment of mitochondria before and after rupturing of the OMM by incubation in the hypotonic buffer revealed that the endogenous ArmC1 was digested by the protease already in intact mitochondria, similar to the OMM protein Tom20 (Fig 2B). We next tested if ArmC1 can be extracted from mitochondria using the sodium carbonate buffer, to address the membrane integration of this protein. Carbonate extraction showed that already at the pH 10.8 the majority of ArmC1 was found in the supernatant and not in the membrane pellet, comparable to the soluble mitochondrial protein Hsp60, and contrary to the OMM anchored protein Tom20 (Fig 2C). Sequence analysis of ArmC1 revealed the presence of conserved armadillo repeat and highly conserved C-terminal segment, but no transmembrane domain (Fig 1D). Taken together, these data demonstrate that ArmC1 is localized both in mitochondria and in the cytosol, and that it is not integrated into the OMM but associates with it, possibly through its C-terminal part.

**Fig 2.**
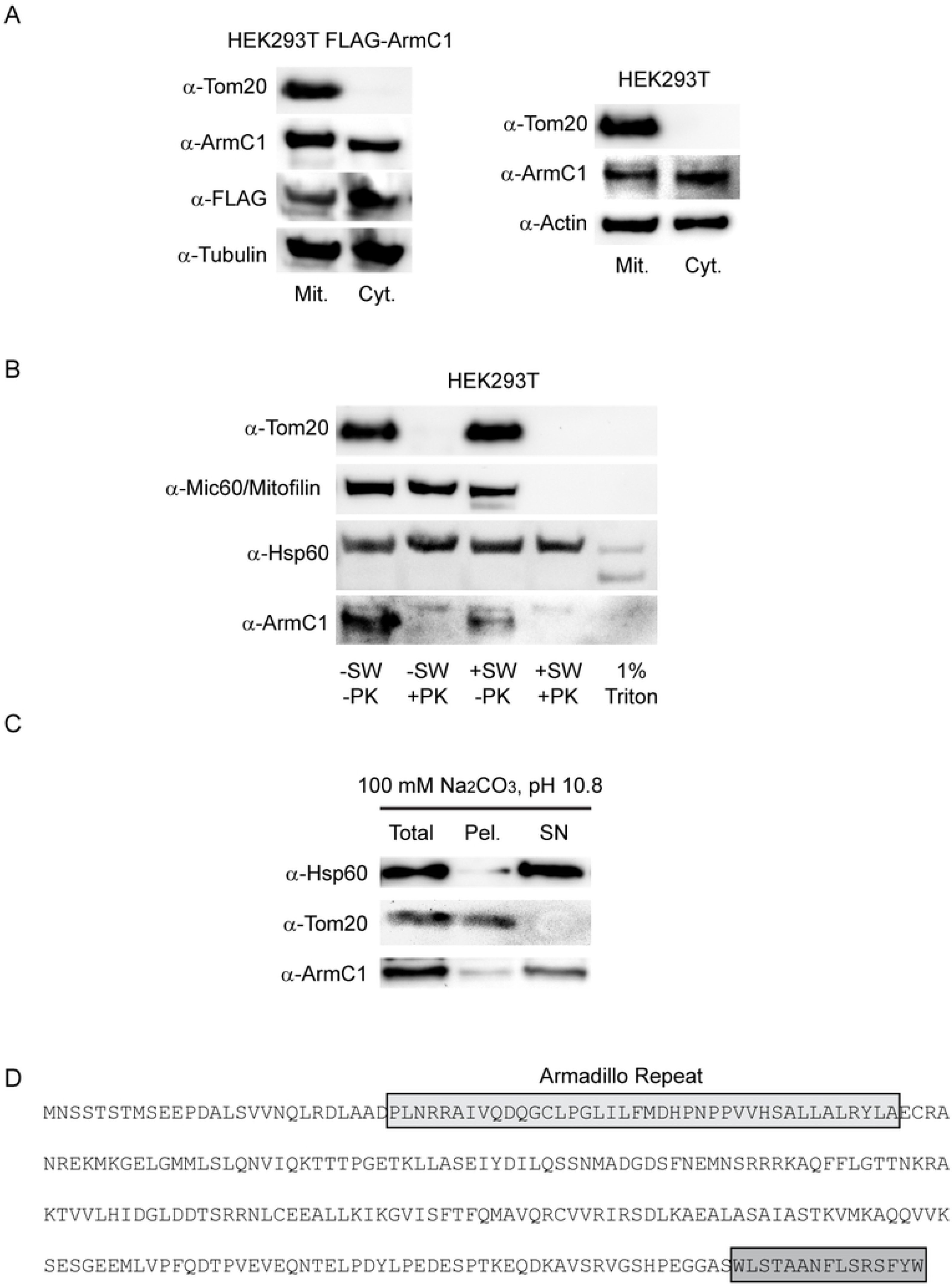
Analysis of mitochondrial association of endogenous ArmC1. (A) HEK293T cells were transfected with pCDNA3 construct encoding for N-terminally FLAG-tagged ArmC1 (FLAG-ArmC1). Transfected and non-transfected HEK293T cells were fractionated into mitochondrial and cytosolic fraction and analyzed by SDS-PAGE and western blot, using antibodies against Tom20, ArmC1, FLAG, tubulin and actin. (B) Mitochondria were isolated from HEK293T cells and subjected to swelling in hypotonic buffer or lysis with 1 % Triton X-100, followed by the treatment with 50 µg/ml protease K. Samples were analyzed by SDS-PAGE and western blot, using antibodies against Tom20, Mic60/Mitofilin, Hsp60 and ArmC1. (C) Mitochondria as in B were subjected to carbonate extraction in 100 mM Na_2_CO_3_, pH 10.8. Total mitochondria, membrane pellet and cytosolic fraction were analyzed by SDS-PAGE and western blot using antibodies against Hsp60, Tom20 and ArmC1. (D) Amino acid sequence of ArmC1 with predicted armadillo repeat and the conserved C-terminal domain in gray boxes. Tom20 – translocase of the outer membrane 20, Hsp60 – heat shock protein 60.

### ArmC1 is a component of the MICOS/MIB complex

ArmC1 has been found to interact with the constituents of the MICOS/MIB complex [17]. The analysis of the protein content of Sam50- and Mic60/Mitofilin-depleted mitochondria also indicated reduction in the amount of ArmC1 when Mic60/Mitofilin, but not Sam50, was downregulated ([14, 19] and not shown). We performed western blot analysis of mitochondria isolated from cell lines where different MICOS/MIB complex subunits were depleted with the help of doxycycline-inducible shRNA-mediated knockdown [10]. We observed that the levels of ArmC1 were strongly reduced upon depletion of Mic60/Mitofilin, as expected, and only mildly affected by the lack of Sam50. Depletion of Mic19/CHCHD3, Mic25/CHCHD6 and Mic23/ApoO did not affect the levels of ArmC1 (Fig 3).

**Fig 3.**
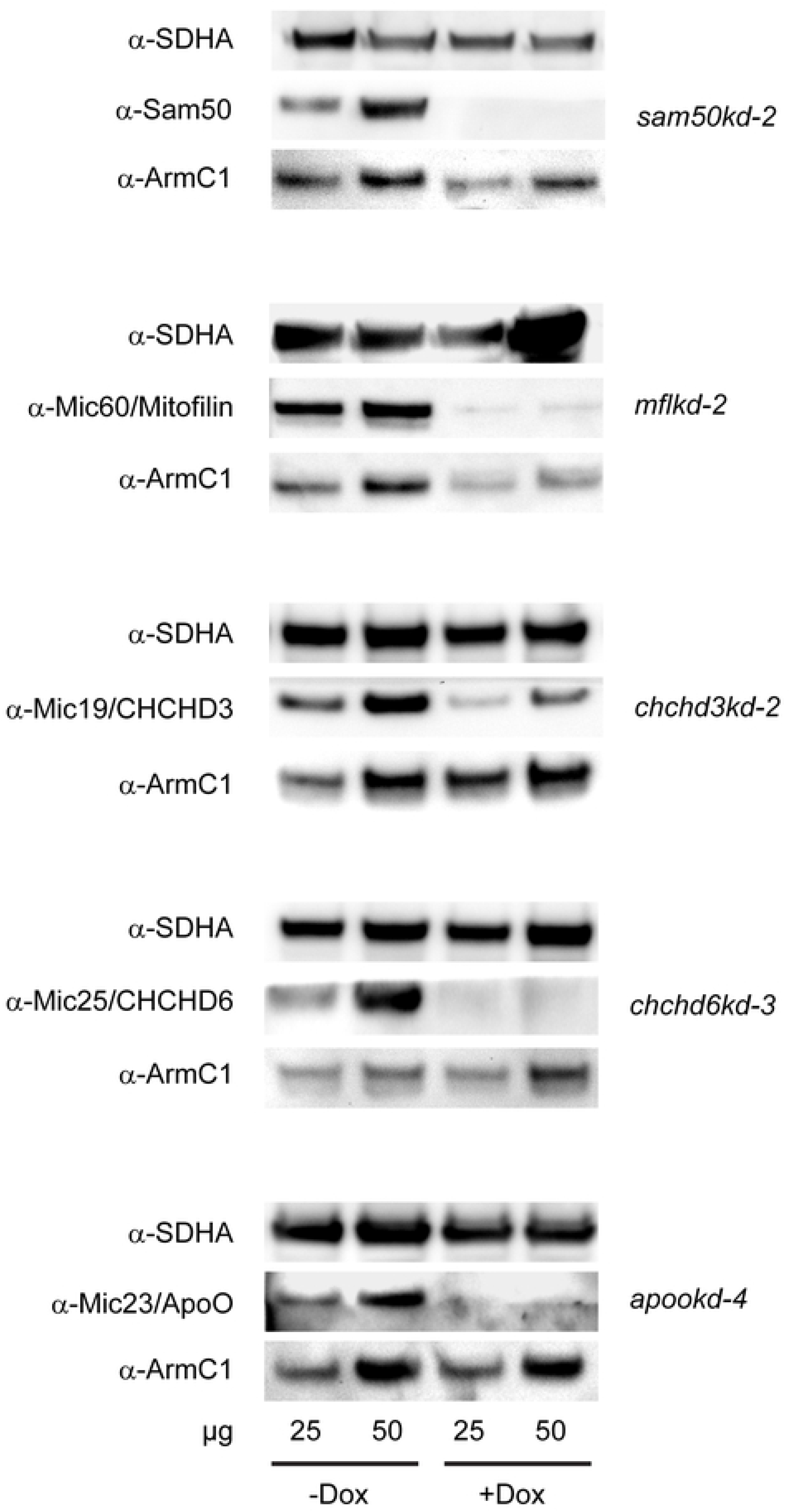
Levels of ArmC1 in the mitochondria of the cells depleted of the MICOS/MIB complex components. Mitochondria were isolated from the cells where a shRNA-mediated knockdown of specific proteins was induced by the addition of doxycycline (Dox) for seven days. 25 and 50 µg of mitochondrial protein was analyzed by SDS-PAGE and western blot using antibodies against SDHA, Sam50, Mic60/Mitofilin, Mic19/CHCHD3, Mic25/CHCHD6, Mic23/ApoO and ArmC1. SDHA, Succinate dehydrogenase complex subunit A, Sam50, sorting and assembly machinery 50, CHCHD, coiled-coil-helix-coiled-coil-helix domain, ApoO, Apolipoprotein O.

To determine the interacting partners of ArmC1, we purified mitochondria from HEK293T cells expressing FLAG-ArmC1 and performed immunoprecipitation (IP) using FLAG antibody. After mass spectrometry analysis of the proteins that were present in the IP from FLAG-ArmC1-containing mitochondria and not in the IP from the control sample, we identified several putative interactors of ArmC1. These included many components of the MICOS/MIB complex, such as Mic60/Mitofilin, DnaJC11 and Mic19/CHCHD3, but also the heat shock binding immunoglobulin protein (BiP) known as Grp78, heat shock cognate 71 kDa protein (Hsp7C), mitochondrial stress-70 protein (Grp75), ATPase family AAA domain-containing protein 3A (Atd3A), and mitochondrial ATP synthase subunit alpha (ATP5F1A/F1α) (Fig 4A). To confirm the IP data, we performed western blot analysis of the IP samples and found that FLAG-ArmC1 indeed co-precipitated with members of the MICOS/MIB complex Mic60/Mitofilin, Mic10/MINOS1, Sam50, and Metaxin 1, but not with Actin or Hsp60 (Fig 4B). Blue-native (BN)-PAGE analysis of protein complexes formed by ArmC1 showed that the protein was mostly present in a high molecular weight protein complex, similar to the one formed by Mic60/Mitofilin and Sam50 (Fig 4C). We conclude that ArmC1 behaves as a component of the MICOS/MIB complex, because it interacts with other subunits of the same complex and is affected by its depletion through downregulation of the central subunit Mic60/Mitofilin.

**Fig 4.**
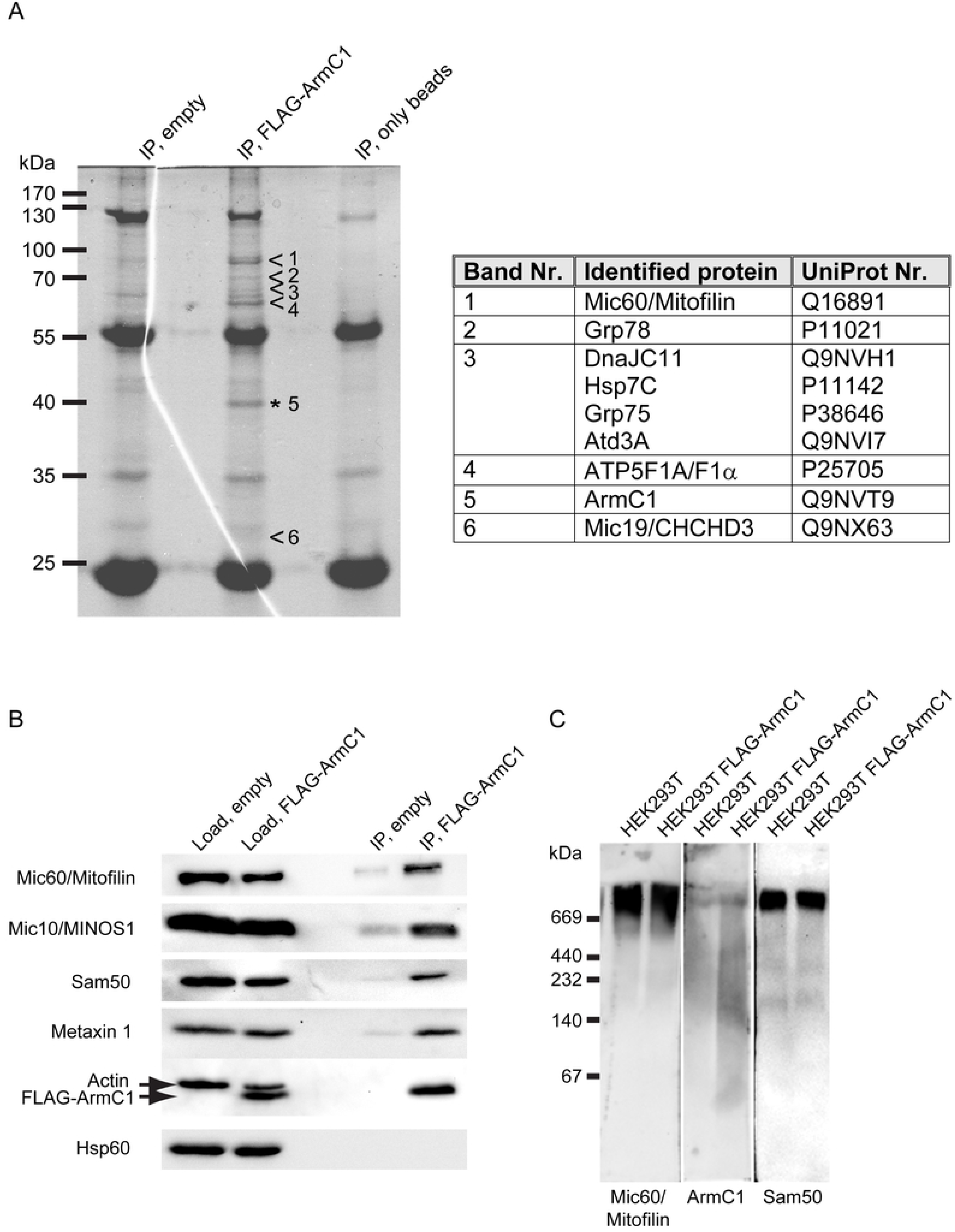
Immunoprecipitation and BN-PAGE analysis of ArmC1. (A) Mitochondria were isolated from HEK293T cells transfected with an empty pCDNA3 plasmid or with the FLAG-ArmC1 pCDNA3 construct, lysed in buffer containing 0.5 % digitonin and 1 mM PMSF, and incubated with Anti-FLAG M2 affinity gel. Eluted proteins were separated using SDS-PAGE and after colloidal coomassie G-250 staining specific bands were excised from the FLAG-ArmC1 sample and analyzed by mass spectrometry in parallel with the respective regions of the gel from the control empty vector sample. Proteins detected in the FLAG-ArmC1 but not in the control sample are listed in the adjacent table. (B) Immunoprecipitation was performed as in A and samples were analyzed by SDS-PAGE and western blot, using antibodies against Mic60/Mitofilin, Mic10/MINOS1, Sam50, Metaxin 1, Actin, FLAG and Hsp60. (C) Mitochondria were isolated from non-transfected HEK293T cells and cells where FLAG-ArmC1 has been expressed with the help of transient transfection of the FLAG-ArmC1 pCDNA3 plasmid. After solubilization with 1% digitonin buffer, samples were analyzed by BN-PAGE and western blot, using antibodies against Mic60/Mitofilin, ArmC1 and Sam50. MINOS1 - mitochondrial inner membrane organizing system 1, Sam50 - sorting and assembly machinery 50, Hsp60 – heat shock protein 60.

### Knockout of ArmC1 leads to mitochondrial fragmentation, but does not influence cristae morphology

We next generated knockout cell lines for ArmC1 in HeLa background using CRISPR/Cas9 technology. Two knockout clones (ArmC1ko-cl. 11 and ArmC1ko-cl. 13) were confirmed for ArmC1 knockout by western blot (Fig 5A and not shown), as well as by sequencing of the corresponding genomic region. ArmC1 knockout did not affect either the cell growth, or the levels of different mitochondrial proteins we tested (Fig 5A and not shown). We performed Seahorse XF Cell Mito Stress Test on ArmC1ko-cl. 11 cells, but again saw no difference in comparison to wild type HeLa cells (Fig 5B). Finally, we analyzed the MICOS/MIB complex by blue native polyacrylamide gel electrophoresis (BN-PAGE) and western blot, but did not observe any major effect of ArmC1 knockout on the appearance of this protein complex (Fig 5C).

**Fig 5.**
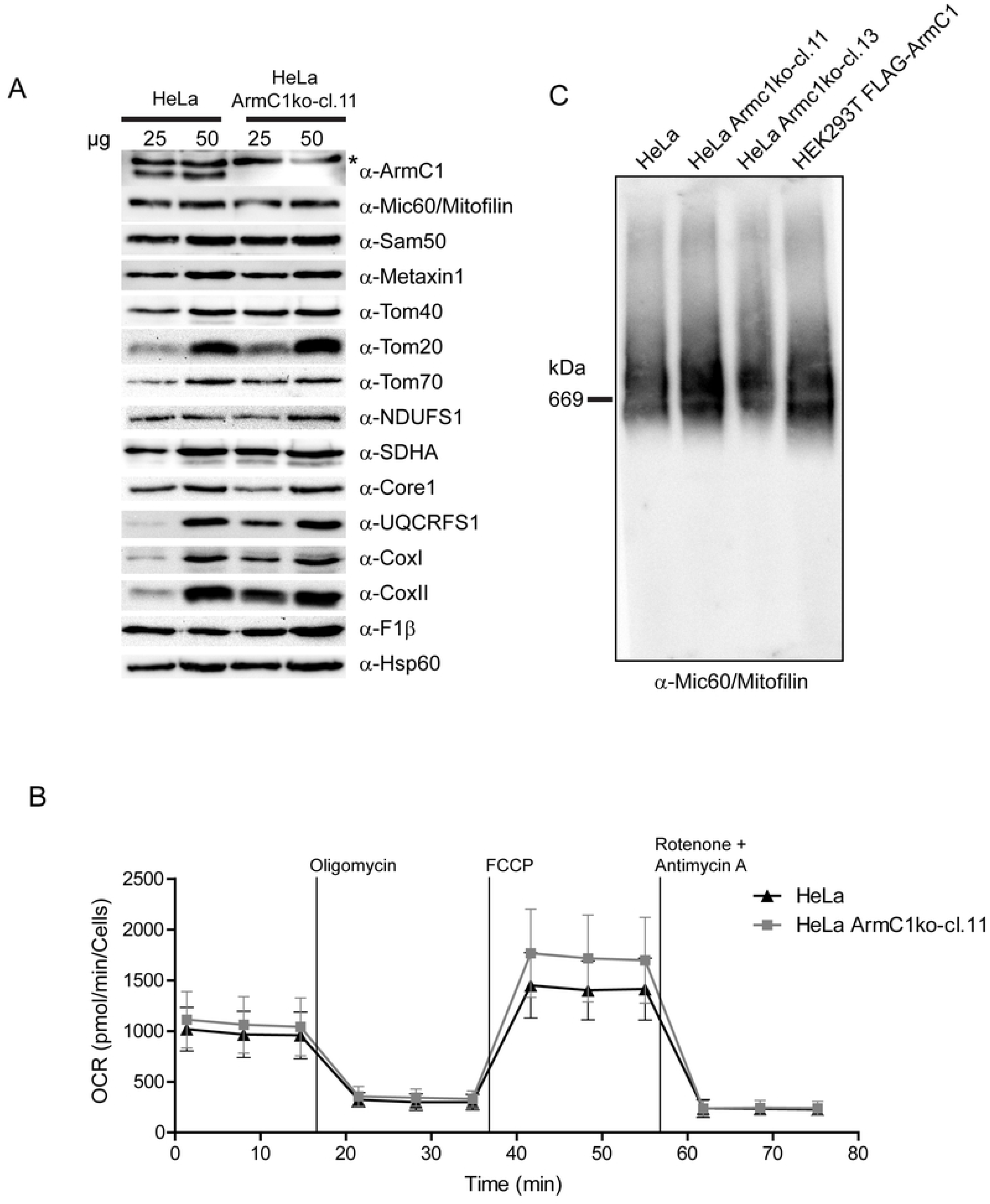
Assessment of mitochondrial protein levels, respiration and MICOS/MIB complex integrity in ArmC1 knockout cells. (A) Mitochondria were isolated from wild type and ArmC1 knockout HeLa cells and 25 and 50 µg of mitochondrial protein was analyzed by SDS-PAGE and western blot with designated antibodies. Asterisk indicates a non-specific band. (B) Oxygen consumption rate (OCR) of wildtype and ArmC1 knockout cells was analyzed by Seahorse Flux Analyzer. Basal respiration, the ATP production, the maximal respiration and the non-mitochondrial respiration were determined by sequentially injecting 2 μM oligomycin, 1 μM FCCP, 0.5 μM rotenone and antimycin A. (C) Mitochondria from wildtype HeLa cells, two different knockdown cell lines of ArmC1 and HEK293T cells transfected with FLAG-ArmC1 pCDNA3 construct were isolated and analyzed by BN-PAGE and western blot, using antibodies against Mi60/Mitofilin, ArmC1 and Sam50. Sam50, sorting and assembly machinery 50, Tom, translocase of the outer mitochondrial membrane, NDUFS1, NADH dehydrogenase [ubiquinone] iron-sulfur protein 1, SDHA, Succinate dehydrogenase complex subunit A, UQCRFS1, Ubiquinol-cytochrome c reductase iron-sulfur subunit 1, Cox, cytochrome oxidase, F1β, ATP synthase subunit beta, Hsp60, heat shock protein 60.

Transmission electron microscopy (TEM) analysis of mitochondrial ultrastructure in wild type HeLa cells versus the ArmC1 knockout cell lines showed no significant difference in cristae morphology (Fig 6A). When we analyzed mitochondrial length in control HeLa cells and cells lacking ArmC1, we identified significant fragmentation of mitochondria. In some cells we also observed clumping of mitochondria in the perinuclear region (Fig 6B). Live cell imaging and quantification of mitochondrial motility in wild type versus knockout cells revealed that deletion of ArmC1 led to a reduced mitochondrial motility (Fig 6C, S1-S3 Movs). In conclusion, ArmC1 knockout has little influence on mitochondrial function, protein content or cristae structure, but affects mitochondrial length and distribution, as well as mitochondrial motility.

**Fig 6.**
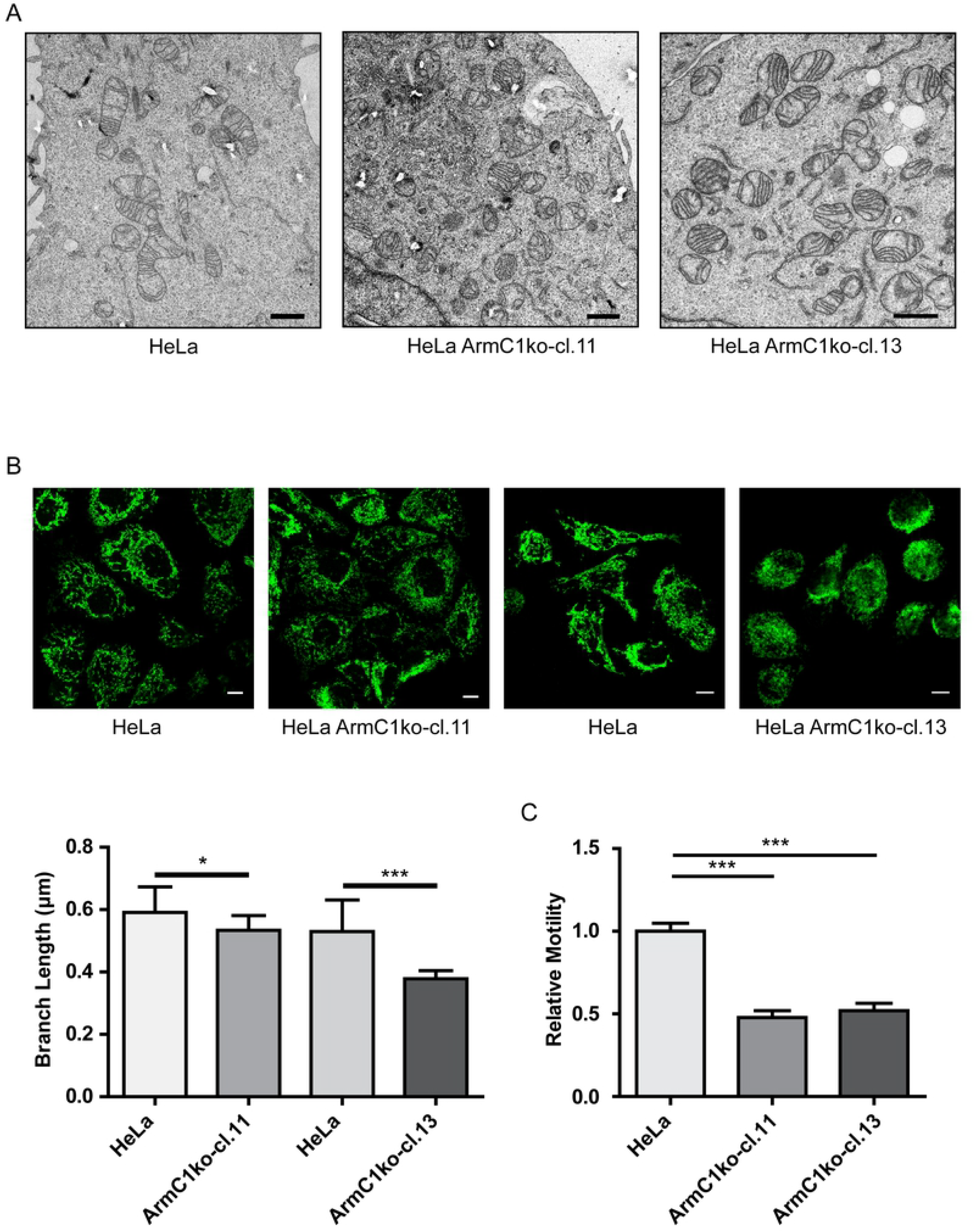
Mitochondrial ultrastructure, morphology and dynamics in ArmC1 knockout cells. (A) Cells of wildtype HeLa and two ArmC1 knockout cell lines were seeded on coverslips, and after fixation with 2.5 % glutaraldehyde prepared and analyzed by transmission electron microscopy. Scale bar is 1 µm. (B) Wildtype HeLa and two ArmC1 knockout cell lines were seeded on coverslips, fixed and stained with antibodies against translocase of the outer membrane (Tom20) protein and secondary antibodies coupled to the Cy2 fluorophore. Pictures were made using confocal microscopy and average mitochondrial branch length was calculated from 10 random fields of view, each containing at least 5 cells. Scale bar is 10 µm. The graph represents mean values ±SD. Significance was calculated using Student’s t test.: * - p≤0.05, *** - p≤0.001. (C) Cells as in A and B were transfected with the pCDNA3 plasmid carrying information for the mitochondrial matrix targeted GFP (CoxVa presequence-GFP), stained with SIR-tubulin and imaged once per minute by confocal microscope for 10-15 minutes. Mitochondrial motility between the frames was measured and normalized to wildtype HeLa results. The graph represents mean values ±SD. Significance was calculated using Student’s t test.: *** - p≤0.001.

## Discussion

In this study, we have characterized ArmC1, a novel putative component of the MICOS/MIB complex in mitochondria. This protein belongs to the family of Armadillo (Arm)-repeat proteins, an evolutionary ancient group present in animals, plants and fungi. Arm-repeat proteins exhibit a common structure of the Arm-repeat domain, although not always sequence similarity [20]. ArmC1, which contains one 42 aa long Arm repeat, has homologues throughout the animal kingdom, and is highly conserved among mammals (>97 % of identity).

The prototypical Arm-repeat protein is β-catenin, an adhesion and signaling protein that interacts with cadherin at adherence junction, but is also present in the cytosol and nucleus, where it is involved in the transduction of Wnt signals [21]. Another example is importin-α, a protein conserved across eukaryotic kingdom, which regulates the transport of proteins into nucleus [20]. Both proteins contain several Arm repeats. Similar to these Arm-repeat proteins, ArmC1 localizes to multiple cellular compartments and can be found both in the cytosol and mitochondria (Fig 1 and 2). All so far characterized components of the MICOS/MIB complex, with an exception of disrupted in schizophrenia 1 (DISC1), are exclusively localized to mitochondria. DISC1, which interacts with Mic60/Mitofilin [22] and has recently been shown to be an IMM component of the MICOS/MIB complex [23], is present in several isoforms, of which two can be found in mitochondria, whereas there is evidence that DISC1 also localizes to cytosol, actin filaments and nucleus [24].

Photobleaching experiments we performed indicated that cytosolic ArmC1 is present as a soluble protein (not shown). Mitochondrial ArmC1 associates, but does not integrate into the OMM (Fig 2). Dual localization of ArmC1 might be connected to its function or there might be a pool of inactive cytosolic ArmC1 that is recruited to mitochondria or shuttled between mitochondria and cytosol upon specific signals, similar to the apoptotic protein Bax [25, 26]. The association of ArmC1 with the mitochondrial surface depends on the C-terminus of the protein. We have identified a 15 aa long conserved region at the end of ArmC1, which might act as a mitochondrial interaction domain, leaving the rest of the protein including the N-terminal Arm-domain exposed to the cytosol.

Data from other groups, as well as our results, demonstrate interactions of mitochondrial ArmC1 with several components of the MICOS/MIB complex [17] (Fig 4). In addition, mitochondrial ArmC1 levels are diminished after the depletion of Mic60/Mitofilin, and ArmC1 is found in the complex of the similar size as the MICOS/MIB, which all corroborates the hypothesis that ArmC1 is a member of the MICOS/MIB complex. The role of ArmC1 in the stability of the MICOS/MIB complex and in mitochondrial function appears, however, to be peripheral, because neither was affected by ArmC1 knockout, and levels of other mitochondrial proteins remained unchanged in the absence of ArmC1 (Fig 5). Considering that the depletion of Sam50 does not have a major effect on the levels of mitochondrial ArmC1 (Fig 3), it is possible that the interaction of ArmC1 with the MICOS complex does not occur through the SAM complex, but directly through the interaction with Mic60/Mitofilin, or maybe through an isoform of DnaJC11, another component of the MICOS/MIB complex with unclear function. In any case, this is a stable interaction, because we were able to co-precipitate ArmC1 with several MICOS/MIB subunits after mitochondria were solubilized with 0.5 % digitonin (Fig 4). What could be the function of ArmC1in mitochondria? Loss of ArmC1 does not have an effect on mitochondrial cristae (Fig 6A), which is to be expected, considering that the MICOS/MIB complex is still intact (Fig 5C). The interaction of ArmC1 with proteins from the heat shock family might be the consequence of the overexpression of the protein, or their requirements for the proper folding of ArmC1 (Fig 4A). We observe, however, that in cells where ArmC1 has been knocked out, mitochondria appear slightly fragmented and their mobility is impaired. This would implicate ArmC1 in the regulation of mitochondrial fission/fusion and/or interaction with cytoskeleton. One of the interactors of ArmC1 we identified was Atd3A, a protein anchored in the IMM that has been shown to interact with the OMM and to play a role in mitochondrial dynamics and stability of mitochondrial nucleoids [27, 28]. Human interactome network results show an interaction of ArmC1 with the mitochondrial fission regulator 1-like (MTFR1L) protein [29], which is predicted to play a role in mitochondrial fission [30]. A recent report identifies a family of Arm repeat-containing proteins encoded by genes located on the X chromosome, the so-called ArmCx cluster, which have evolved from a common ancestral gene ArmC10 and localize to mitochondria [31]. The ancestral ArmC10 protein has been shown to be a substrate of AMP-activated protein kinase (AMPK) and to be involved in mitochondrial fission [32]. ArmC1 is also phosphorylated at threonine 137, serine 189 and serine 260 [33], which might indicate the regulation of its function through phosphorylation. ArmCx3/Alex3 is involved in the regulation of mitochondrial dynamics and trafficking and interacts with the Kinesin/miro/Trak2 complex [31]. Interestingly, the overexpression of GFP-tagged ArmCx3/Alex3 in HEK293AD and 12 DIV hippocampal neurons leads to mitochondrial aggregation [31], which is highly similar to what we observe upon overexpression of GFP-ArmC1 in HeLa cells (Fig 1B). The difference, however, is that all ArmCx proteins possess mitochondrial targeting signals and transmembrane anchors in their N-terminus, and up to six Arm-repeats in their C-terminus, whereas ArmC1 has a targeting sequence in its C-terminus and only one Arm-repeat in its N-terminus and also does not integrate into the OMM. Nevertheless, the similarity between ArmC1 and ArmCx proteins and the resemblance of overexpression phenotypes could point to the common function. Therefore, ArmC1 could represent a link between the MICOS/MIB complex and mitochondrial dynamics and distribution, adding to the functional diversity and importance of this major organizer of mitochondrial cristae.

## Materials and Methods

### Cell lines and cell culture

Human embryonic kidney epithelial (HEK293T) cells (ATCC CRL-11268) and human cervical carcinoma (HeLa) cells (ATCC CCL-227) were obtained from the American Type Culture Collection. *sam50kd-2*, *mflkd-2*, *chchd3kd-2*, *chchd6kd-3* and *apookd-4* [10] were generated from HeLa cells using lentiviral-based shRNA expression system [34]. ArmC1 knockout cell lines were generated using pSpCas9n(BB)-2A-GFP vector and two gRNAs as described [35]. The gRNA sequences are: CGTCAGGCTCTTCACTCATGG and CAAAGCGGAGTGGACGACTGG. HeLa-based cell lines and HEK293T cells were cultivated in RPMI-1640 and DMEM (Gibco/Thermo Fisher scientific, Massachusetts, USA), respectively, supplemented by 10 % fetal bovine serum (FBS) (Sigma/Merck, Darmstadt, Germany) and 1 % Penicillin/Streptomycin (Gibco/Thermo Fisher scientific, Massachusetts, USA) at 37 °C/5 % CO_2_. For the induction of shRNA-mediated knockdown, cells were cultivated for 7 days in the presence of 1 µg/ml doxycycline (BD Biosciences, Franklin Lakes, New Jersey, USA).

### Cloning and transfection

ArmC1 gene was amplified from HeLa cDNA and cloned into the pCDNA3 vector (Thermo Fisher Scientific, Massachusetts, USA) where previously GFP or FLAG sequences were introduced, enabling N- or C-terminal fusion and tagging. HeLa cells were transfected using Viromer® RED (230155; Biozym, Oldendorf, Germany) according to manufacturer’s instructions. HEK293T cells were transfected using calcium phosphate transfection method. For this, cells were grown on 15 cm tissue culture dishes. Calcium phosphate precipitates were generated by mixing 40 µg of plasmid DNA together with CaCl_2_ solution and the 2xHBS buffer in 4 ml volume to a final concentration of 0.25 M CaCl_2_, 25 mM HEPES pH 7.05, 70 mM NaCl and 0.75 mM Na_2_HPO_4_. The mixture was added to cells dropwise to a total volume of 20 ml after addition of 25 µM chloroquine. Medium was changed on the following day and 36 h later mitochondria were isolated from transfected cells.

### Microscopy

For immunostaining cells were seeded on cover slips and after treatment fixed with 4 % PFA. The fixed cells were washed with PBS and then permeabilized and blocked with a buffer containing 10 % goat serum and 0.2 % Triton X-100 in PBS. Cells were afterwards incubated with the primary antibody in antibody-dilution-buffer (3 % goat serum and 0.05 % Tween20 in PBS) for 1 h at RT. After wash and 10 minutes incubation in blocking buffer, samples were incubated with the corresponding secondary antibody diluted in antibody-dilution-buffer for another 50-60 min. In the final step, the cover slips were mounted onto glass-slides using 2.5 % Mowiol-DABCO (Carl Roth, Karlsruhe, Germany). Images were acquired on a TCS SP5 confocal microscope (Leica Biosystems, Wetzlar, Germany) using a 63x oil-immersion UV objective with a numerical aperture of 1.4 or, for superresolution imaging, on a Zeiss (Oberkochen, Germany) ELYRA S.1 SR-SIM structured illumination platform using a Plan-Apochromat 63x oil-immersion objective with a numerical aperture of 1.4. Reconstruction of superresolution images were performed using the ZEN image-processing platform with a SIM module.

### Quantification and statistical analysis

For quantification of mitochondrial length, HeLa wildtype and ArmC1 knockout cells were seeded on coverslips, fixed and stained with Tom20 antibodies, followed by Cy2-coupled secondary antibodies. Images were taken by confocal microscopy on a TCS SP5 confocal microscope (Leica Biosystems, Wetzlar, Germany) using a 63x oil immersion UV objective with a numerical aperture of 1.4 at 1024 x 1024 px resolution. For image analysis, length of mitochondrial fragments from at least ten random different fields of view (five and more cells per field of view) was measured as previously described [36].

For assessment of mitochondrial motility, human Hela cells, wildtype and ArmC1 knockout cells were seeded in 35 mm high glass bottom dishes (81158; Ibidi, Martinsried, Germany). The following day, the cells were transfected using Viromer® RED (230155; Biozym, Oldendorf, Germany) with a pCDNA3 plasmid containing GFP fused to the presequence of CoxVa to visualize mitochondria. 24 h post transfection the cells were incubated in SIR-tubulin (SC002; Spirochrome, Stein am Rhein, Germany). After washing, the medium was changed to a 37 °C warm RPMI-1640 containing 25 mM Hepes (without phenol red and Ca^2+^/Mg^2+^). The samples were imaged on a TCS SP5 confocal microscope (Leica Biosystems) using a 63x oil immersion UV objective with a numerical aperture of 1.4, at 37 °C. The images were recorded once per minute with the Leica Biosystems Application Suite at 1024 x 1024 px resolution. For image analysis we used a modified version of the ImageJ macro MitoCrwler [36]. Shortly, we opened the movies with Fiji/ImageJ and extracted the GFP channel. Background was subtracted by the rolling ball algorithm (diameter: 50). Changes in the GFP distribution due to mitochondrial motility between two adjacent frames resulted in difference of a thresholded binary image. Therefore, the macro measured the amount of difference between subsequent time frames by the “Analyze Particles” function of ImageJ.

Statistical calculations were performed using GraphPad Prism 6.0. Statistical significance was calculated using Student’s t test.

### Mitochondrial isolation and fractionation

Isolation of mitochondria from cells was performed as described previously [37]. Carbonate extraction, cellular fractionation, and opening of the OMM by swelling were performed as already described [38, 39]. The oxygen consumption rate (OCR) was measured using the Seahorse Bioscience XFe96 Extracellular Flux Analyzer (Seahorse Bioscience, Agilent, Santa Clara, California, USA) according to manufacturer’s protocol [40]. In brief, 2×10^4^ cells were seeded in Seahorse 96 well XF Cell Culture Microplate in a total volume of 80 μl XF Base medium supplemented with 1 mM pyruvate, 2 mM L-glutamine and 10 mM glucose (pH 7.4 ± 0.05). The basal respiration, the ATP production, the maximal respiration and the non-mitochondrial respiration were determined by sequentially injecting 2 μM oligomycin, 1 μM FCCP, 0.5 μM rotenone and antimycin A, respectively. Data were normalized to the cell number of a replicate plate, which was quantified by crystal violet staining.

### Protein analysis

SDS-PAGE, western blot and BN-PAGE were performed as described previously [19]. For BN-PAGE samples were lysed in 1 % digitonin buffer (1 % digitonin (Sigma/Merck, Darmstadt, Germany) in 20 mM Tris–HCl, 0.1 mM EDTA, 1 mM PMSF, 50 mM NaCl, 10 % (v/v) glycerol, pH 7.4) and analyzed on 4-10 % or 4-16.5 % polyacrylamide gradient gels [41]. Immunoprecipitation was performed using Anti-FLAG M2 affiinity gel (Sigma/Merck, Darmstadt, Germany) according to manufacturer’s protocol. Lysis buffer was supplemented with 1 mM Phenylmethylsulfonylfluorid (PMSF) and 0.5 % digitonin (Sigma/Merck, Darmstadt, Germany). Binding of the proteins was performed over night at 4 °C and the proteins were eluted from the beads using Laemmli sample buffer. Gels were stained using colloidal coomassie G-250, and bands appearing in FLAG-ArmC1, but not the control sample were excised from the gel. At the same time, respective areas of the gel lanes of the control samples were excised, too. All gel fragments were analyzed by mass spectrometry as described [19].

### Antibodies

FLAG, Actin and Mic23/ApoO antibodies were purchased from Sigma (St. Louis, Missouri USA), Tom20 and F1β from BD Transduction Laboratories (Franklin Lakes, New Jersey, USA), Mic60/Mitofilin, Mic19/CHCHD3 and Mic25/CHCHD6 from Abcam (Cambridge, UK), SDHA, Core1 and CoxI from Invitrogen/Thermo Fisher Scientific (Massachusetts, USA), Mic10/MINOS1, UQCRFS1 and CoxII from GeneTex (Irvine, California, USA), Tubulin, Tom40 and NDUFS1 from Santa Cruz (Dallas, Texas, USA), Hsp60 from Enzo Life Sciences (Farmingdale, New York, USA) and ArmC1 from Life Technologies (Carlsbad, California, USA). Sam50, Metaxin 1 and ArmC1 antibodies were raised in rabbits against the full-length 10xHis-tagged protein. Rat Tom70 antibody (crossreactive with human Tom70) was a kind gift from K. Mihara. Secondary anti-rabbit and anti-mouse antibodies coupled to Cy3 or Cy5 fluorophores were purchased from Dianova (Hamburg, Germany).

## Acknowledgments

We thank A. Schulze and S. Janaki-Raman for the help with Seahorse, and C. Stigloher, D. Bunsen and C. Gehrig from the Imaging Core Facility, University of Würzburg for the help with electron microscopy.

**S1 Mov. Live cell imaging of HeLa WT cells.** HeLa cells were transfected with a pCDNA3 plasmid containing information for mitochondria-targeted GFP. 24 h post transfection microtubules were stained using SIR-tubulin. The samples were imaged using confocal microscope, with the images recorded once per minute.

**S2 Mov. Live cell imaging of HeLa ArmC1-ko cl.11 cells.** ArmC1-ko cl.11 cells were transfected with a pCDNA3 plasmid containing gene for mitochondria-targeted GFP for mitochondria visualization. 24 h later microtubules were stained using SIR-tubulin. The samples were imaged using confocal microscope, with the images recorded once per minute.

**S3 Mov. Live cell imaging of HeLa ArmC1-ko cl.13 cells.** ArmC1-ko cl.13 cells were prepared for live cell imaging by transfection with a pCDNA3 plasmid containing gene for mitochondria-targeted GFP for mitochondria visualization and 24 h later by staining microtubules using SIR-tubulin. The samples were imaged using confocal microscope, with the images recorded once per minute.

